# Vitamin D3 ameliorates R-loop-induced replication stress and chromosomal instability in MED12-mutant uterine fibroids

**DOI:** 10.1101/2025.09.17.676564

**Authors:** Sribalasubashini Muralimanoharan, Ana Corachán, Azad Khosh, Sierra Hathaway, Susivarshini Karthigayan, Claire Schenken, Nicholas Stansbury, Robert Schenken, Maria Victoria Bariani, Qiwei Yang, Eloise Dray, Mazhar Adli, Hortensia Ferrero, Ayman Al-Hendy, Thomas G. Boyer

## Abstract

Uterine fibroids (UFs) are the most important benign neoplastic threat to women’s health worldwide, with no long-term noninvasive treatment options currently available. Among known UF driver alterations, somatic mutations in Mediator subunit MED12 are by the far the most prevalent, accounting for up to 80% of these clinically significant lesions. Although it is presently unclear how MED12 mutations trigger neoplastic transformation, MED12-mutant UFs are nonetheless characterized by significant chromosomal loss and rearrangement, suggesting genomic instability as a driving force in tumor development. However, the basis by which MED12 mutations drive genomic instability is not known. Herein, we show that R-loop-driven replication stress in MED12-mutant UFs leads to DNA under-replication and mitotic segregation errors that drive chromosomal instability. Notably, we find that vitamin D3 (VD3), a modifiable risk factor in UF development, suppresses pathogenic R-loop accrual and ameliorates replication stress-driven chromosomal instability, contributing to growth inhibition of patient-derived MED12-mutant UF xenografts in vivo. Altogether these findings uncover a molecular basis by which the predominant UF driver converges with a known risk factor at the interface of genomic instability, with significant translational implications for personalized UF prevention and treatment.

## Introduction

Uterine fibroids (UFs; leiomyomas) are benign monoclonal neoplasms of the myometrium (MM) and represent the most common non-cutaneous tumors in women worldwide (1). Tumors occur in ∼77% of women overall and are clinically manifest in ∼25% by age 45 (2). Although benign, these tumors are nonetheless associated with significant morbidity; they are the primary indication for hysterectomy, and a major source of gynecologic and reproductive dysfunction, ranging from abnormal uterine bleeding and pelvic pain to infertility, recurrent miscarriage, and pre-term labor (1, 2). Accordingly, the annual US health care costs associated with UFs have been estimated at ∼$42.2 billion (3).

Despite their high prevalence and significant morbidity, current treatment options for UFs are limited. Curative surgery (hysterectomy) destroys reproductive function, while uterine-sparing surgical and radiologic alternatives are variously associated with reduced long-term reproductive function and/or high tumor recurrence rates (4, 5). On the other hand, current hormonal therapies are contraindicated in women actively pursuing a pregnancy, and otherwise effective only during use, which is limited due to long-term safety and other concerns (4–6). These limitations highlight a pressing need for the development of novel safe, efficacious, and fertility-compatible options for UF prevention and treatment.

We and others have previously reported that an inverse correlative association exists between circulating vitamin D (VD3) levels and patient UF risk (7–9), initially suggesting that VD3 supplementation might represent a potential preventive and/or therapeutic option against UFs. Indeed, this hypothesis is now supported by a significant body of preclinical data demonstrating that VD3 can suppress UF cell and tumor growth through proven antiproliferative and antifibrotic properties in vitro and/or in vivo (10–12). More recently, clinical studies have revealed that VD3 treatment can restrict or even reduce tumor growth in women bearing UFs (11, 13–15). These findings are particularly relevant for African American (AA) women, who are disproportionately prone to increased UF risk and hypovitaminosis D. Thus, compared to European American (EA) women, AA women experience a 3-fold higher incidence and relative risk of UF disease, as well as earlier disease onset and more profound symptomatic disease (1, 16). Furthermore, AA women are 10-times more likely to be VD3-deficient than EA women (17–19). However, our current understanding of the relationship between VD3 and UF pathogenesis at the molecular and biochemical levels remains incomplete.

The prevailing model for UF pathogenesis invokes the genetic transformation of a single MM stem cell into a tumor-initiating cell that seeds and sustains clonal tumor growth, characterized by increased cell proliferation and abundant ECM production, under the influence of endocrine, autocrine, and paracrine growth factor and steroid hormone receptor signaling (1, 20–23). The genetic drivers dominantly responsible for cell transformation have now been largely identified. Among these, somatic exon 1/2 mutations in the gene encoding the RNA polymerase II Mediator subunit MED12 are by far the most prevalent, accounting for 50-80% of UFs in women of diverse racial and ethnic origins (24–29). Notably, recent work suggests that the MED12 mutation rate may be higher in UFs from AA compared to EA or Asian women (30–33). At the biochemical level, all UF-driver mutations in MED12 are known to disrupt its ability to activate CycC-CDK8 kinase in Mediator, leading to overall loss of Mediator kinase activity (24, 29, 34–37). Although it is presently unclear how MED12 mutations and loss of Mediator kinase activity precipitate neoplastic transformation, MED12-mutant UFs are nonetheless characterized by significant chromosomal loss and rearrangement, suggesting genomic instability as a driving force in tumor development (28, 38–41). For example, prior studies have revealed that over 60% of UFs with an abnormal karyotype carry MED12 mutations (27, 28). Furthermore, UF tumors arising in MED12-mutant mice were found to carry chromosomal aberrations, many with syntenic counterparts on human chromosomes, including 1p, 1q, 2q, 6p21, and 18p, known to be rearranged in human UFs (40).

Of note, some of these genome-wide alterations included complex chromosomal rearrangements resembling chromothripsis which has also been observed in human MED12-mutant UFs (38, 42). Nonetheless, the mechanistic basis by which MED12 mutations drive genomic instability and tumor development remains unclear.

In this regard, we recently showed that MED12 mutation-positive UFs, compared to patient-matched MED12 mutation-negative UFs and MM, accrue pathogenic R-loops and replication stress (43). Further, we found that these phenotypes could be recapitulated and functionally linked in uterine smooth muscle cells (UtSMCs) by chemical inhibition of Mediator-associated CycC-CDK8 kinase activity that is disrupted by UF driver mutations in MED12 (43). Altogether, these findings reveal that MED12-mutant UFs are uniquely prone to R-loop-induced replication stress, uncovering a potential therapeutic vulnerability in this dominant UF subclass. Nonetheless, whether and how replication stress occurring in MED12-mutant UFs contributes to chromosomal instability and UF development has not heretofore been investigated. Further, whether and how factors that drive UF formation, including MED12 mutations and chromosomal instability, are biochemically linked with established factors of disease risk, including VD3 and race, is largely unknown. Herein, we provide a mechanistic basis to couple UF etiology and established risk factors through a functional interplay between VD3 and R-loop-dependent replication stress in MED12-mutant UFs. We show that R-loop-driven replication stress in MED12-mutant UFs leads to DNA under-replication in mitosis and hallmarks of genomic instability, including chromosomal segregation defects and micronuclei formation. Notably, we find that VD3 treatment, by suppression of pathogenic R-loop accrual, ameliorates replication stress-driven chromosomal instability, leading to inhibition of MED12-mutant UF tumor xenografts in vivo. Furthermore, we provide preclinical proof-of-concept for treatment of human UFs with the FDA-approved nonhormonal VD2 analog Doxercalciferol (DCL). Altogether these findings uncover a molecular basis by which mutant MED12 and VD3 converge at the interface of genomic instability, with significant translational implications for personalized UF prevention and treatment.

## Results

### VD3 ameliorates R-loop-dependent replication stress in MED12-mutant UFs

Previously, we showed that MED12-mutant UFs are characterized by aberrant accrual of R-loops and altered replication fork dynamics indicative of replication stress (43). Herein, we confirmed and functionally linked these phenotypes in primary cells from patient-matched MM and MED12-mutant UFs (Table S1). First, immunocytochemical analysis using the RNA-DNA hybrid-specific antibody S9.6 revealed significantly elevated levels of R-loops in primary cells from MED12-mutant UFs compared to those from patient-matched MM (**Fig. 1A**). Second, single molecule DNA fiber analysis following IdU/CldU pulse-labeling (**Fig. 1B**) revealed significantly altered replication fork dynamics in MED12-mutant UF cells compared to those from patient-matched MM, including increased and decreased numbers of stalled and restarted replication forks, respectively (**Fig. 1C**), reduced replicative DNA fiber lengths (**Fig. 1D**), and increased numbers of bidirectional asymmetric forks, which derive from asymmetric fork progression through replication fork arrest (**Fig. 1E**). Thus, MED12-mutant UFs are uniquely characterized by both aberrant accrual of R-loops and replication stress. Importantly, these phenotypes were effectively reversed by ectopic overexpression of RNase H1 that degrades RNA-DNA hybrids, indicating that replication stress arising in MED12-mutant UF cells is R-loop-dependent (**Fig. 1A-E**; **Fig. S1A**).

**Figure 1.**
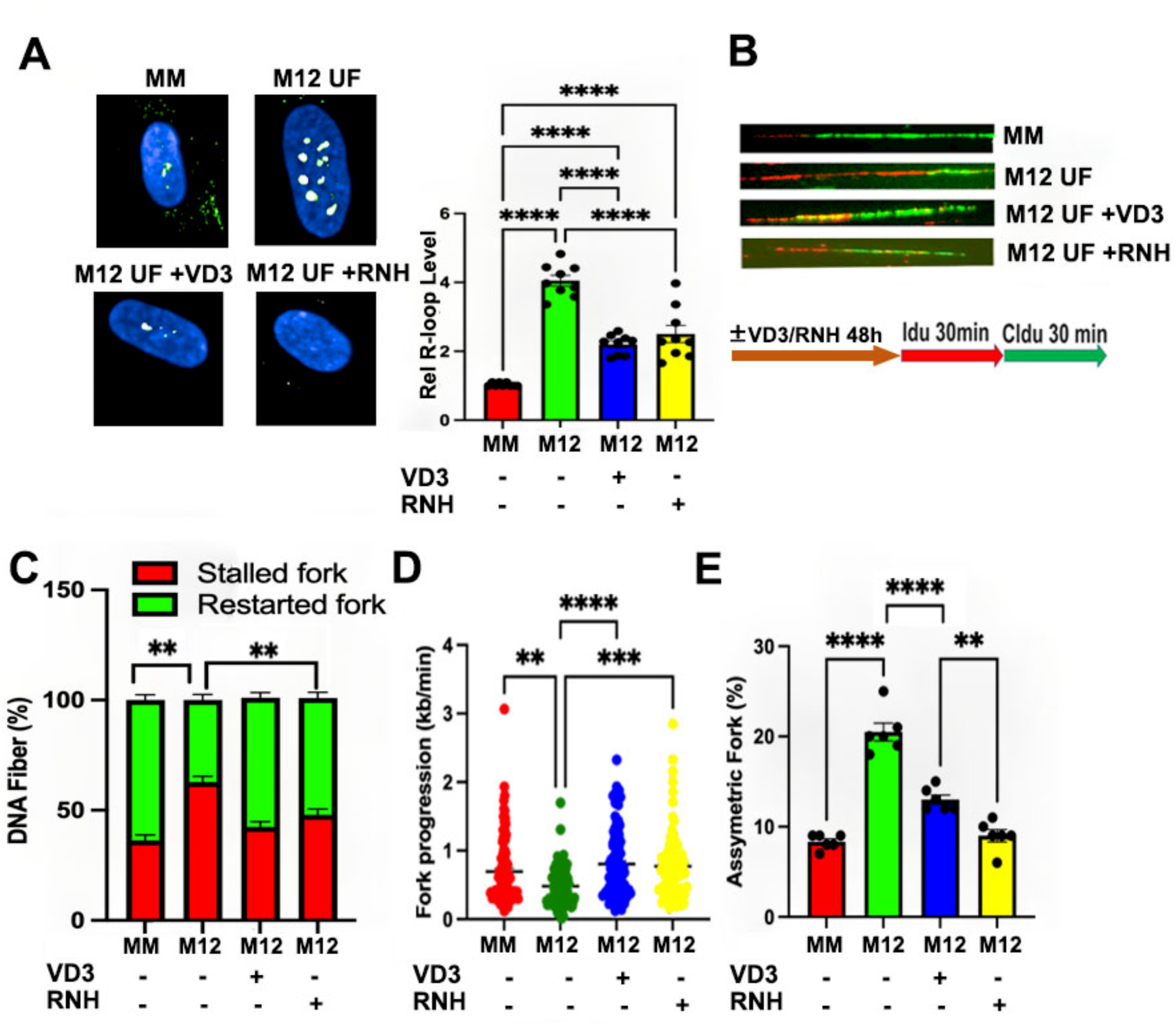
VD3 ameliorates R-loop-dependent replication stress in MED12-mutant UFs. (**A**) Representative images (*left*) and quantification (*right*) of R-loop levels in primary patient-paired MM and MED12-mutant (M12) UF cells by immunocytochemical analysis using the RNA-DNA hybrid-specific antibody S9.6. Where indicated, cells were treated for 48h without (-) or with (+) VD3 (100nM) or transduced with RNase H1 (RNH)-expressing lentivirus prior to R-loop staining. Data are plotted as fold-change in S9.6 antibody signal intensity relative to untreated MM cells. Quantification was determined from a minimum of n=50 cells in each of 3 technical replicates from 3 independent experiments (n≥450 total cells) per treatment condition. (**B**) Representative images of combed DNA fibers from primary MM/UF cells pulse-labeled with thymidine analogues IdU (red) and CldU (green). As indicated, cells were treated without or with VD3 (100nM) or transduced with RNase H1 (RNH)-expressing lentivirus 48h prior to pulse labeling. The pulse labeling scheme is shown below. Replication fork dynamics were measured as described in panels **C**-**E** from a minimum of n=35 fibers in each of 3 technical replicates from 2 independent experiments (n≥210 total fibers) per treatment condition. (**C**) Percent of singly labeled fibers retaining only first (IdU) or second (CldU) label indicative of stalled or new forks, respectively. (**D**) Velocity of fork progression determined from ldU tract lengths (kb/min). (**E**) Percent of bidirectional asymmetric forks among total fork numbers. Significance calculated using One-Way ANOVA and Tukey’s Post hoc test, ****p < 0.0001, ***p<0.001; **p<0.01.

Based on both preclinical and clinical data demonstrating tumor reductive properties associated with VD3 (11, 13–15), we investigated the impact of VD3 on R-loop-induced replication stress in MED12-mutant UFs. Strikingly, we found that VD3 effectively suppressed aberrant R-loop accrual in primary cells from MED12-mutant UFs (**Fig. 1A**) and did so in a dose-dependent manner (**Fig. S2A**). Consistent with its ability to suppress R-loop accumulation in primary MED12-mutant UF cells, we also observed that VD3 treatment ameliorated R-loop-dependent replication fork defects arising in these cells (**Fig. 1B**), effectively decreasing and increasing numbers of stalled and restarted forks, respectively (**Fig. 1C**), restoring replication fork speeds (**Fig. 1D**), and decreasing asymmetrical bidirectional forks (**Fig. 1E**).

Previously, we showed that UF-driver mutations in MED12 all similarly disrupt Mediator-associated CDK8/19 kinase activity, revealing the first and heretofore only known biochemical defect associated with these pathogenic mutations (24, 29, 34–37). In fact, many of the cellular and molecular phenotypes attributed to MED12 UF driver mutations can be recapitulated by direct chemical inhibition of Mediator-associated CDK8/19 kinase activity in either patient-derived MM stem cells or hTERT-immortalized uterine smooth muscle cells (UtSMCs) (43–45), rendering pharmacological suppression of Mediator kinase activity a valuable and experimentally tractable system to model the pathogenic consequences of MED12 mutations. Notably, this phenotypic concordance extends to aberrant accrual of R-loops and R-loop-dependent replication stress, which we previously documented in UtSMCs treated with the highly specific CDK8/19 kinase inhibitor CCT251545 (CCT) (43, 46). Thus, we showed that Mediator kinase inhibition in UtSMCs with CCT led to pathological accrual of R-loops and replication fork defects indicative of replication stress that could be reversed by ectopic overexpression of RNase H1 (43). Herein, we used this system (**Fig. S2B**) to examine the impact of VD3 on both aberrant R-loop accrual and R-loop-induced replication stress in Mediator kinase-inhibited UtSMCs. Consistent with our observations in primary MED12-mutant UF cells, we found that VD3 treatment suppressed both aberrant R-loop accrual (**Fig. 2A; Fig. S2C)** and R-loop dependent replication fork defects (**Fig. 2B**) in Mediator kinase-inhibited UtSMCs, resulting in decreased and increased numbers of stalled and restarted forks, respectively (**Fig. 2C**), restoration of replication fork speeds (**Fig. 2D**), and reduced numbers of asymmetrical bidirectional forks (**Fig. 2E**). Further, we also observed that VD3 treatment suppressed aberrant R-loop accrual in CRISPR-engineered MED12-mutant (G44N) human TERT-immortalized UtSMCs (47) compared to their corresponding MED12 WT counterparts (**Fig. 3A and Fig. S3**).

**Figure 2.**
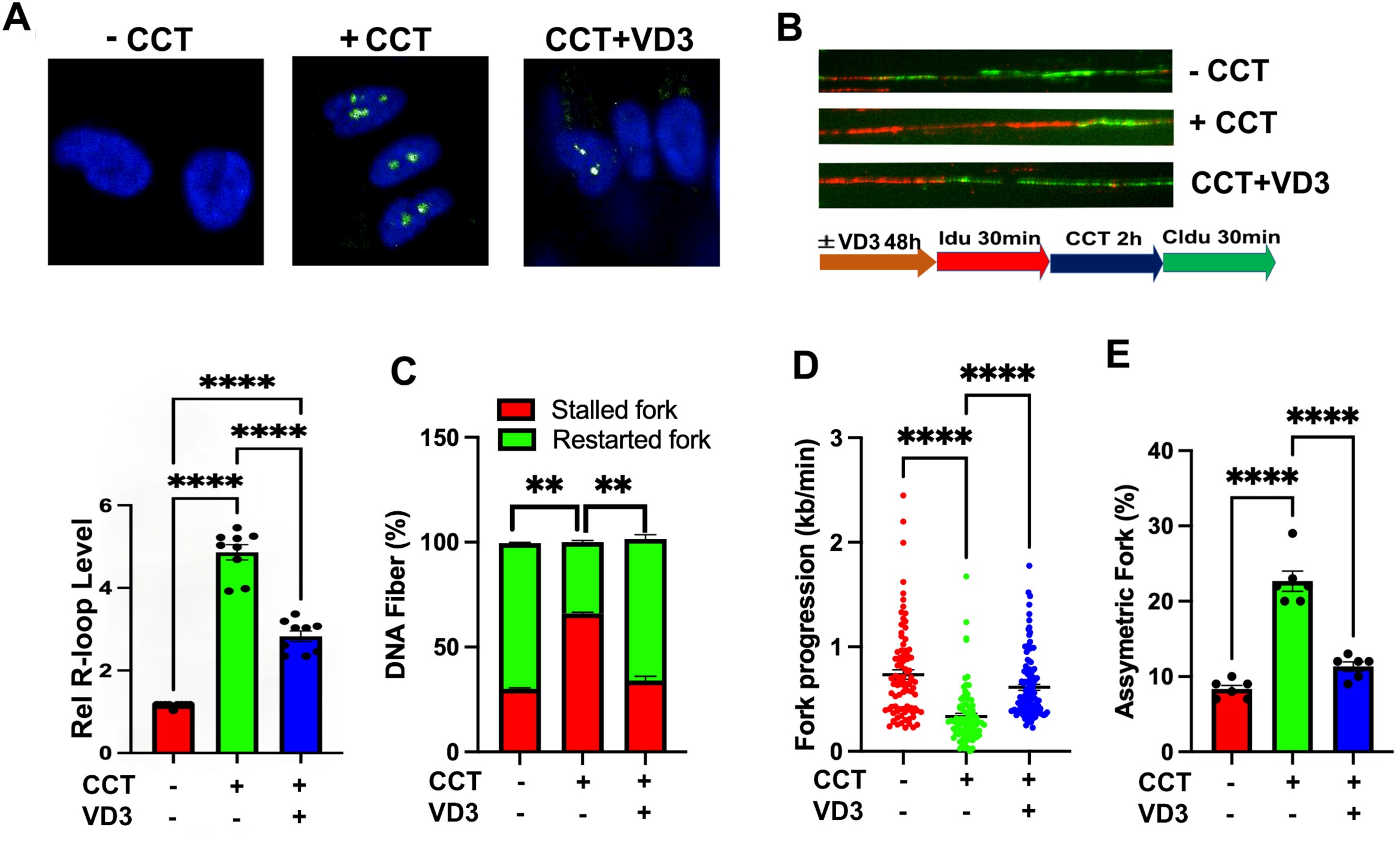
VD3 ameliorates R-loop dependent replication stress in Mediator kinase-inhibited UtSMCs. (**A**) Representative images (*top*) and quantification (*bottom*) of R-loop levels in UtSMCs assessed by immunocytochemical analysis using the RNA-DNA hybrid-specific S9.6 antibody. Where indicated, cells were treated without (-) or with (+) CDK8/19 inhibitor (CCT; 100nM for 24h) and/or VD3 (100nM for 48h) prior to R-loop staining. Data are plotted as fold-change in S9.6 antibody signal intensity relative to untreated UtSMCs. Quantification was determined from a minimum of n=50 cells in each of 3 technical replicates from 3 independent experiments (n≥450 total cells) per treatment condition. (**B**) Representative images of combed DNA fibers from IdU/CldU pulse-labeled UtSMCs cells treated without (-) or with (+) CDK8/19 inhibitor (CCT; 100nM). Where indicated, cells were treated with VD3 (100nM) for 48h prior to pulse labeling. The pulse labeling scheme is shown below. CCT was added for 2h between 30 min Idu (red) and CldU (green) pulses. Replication fork dynamics were measured as described in panels **C**-**E** from a minimum of n=35 fibers in each of 3 technical replicates from 2 independent experiments (n≥210 total fibers) per treatment condition. (**C**) Percent of singly labeled fibers retaining only first (IdU) or second (CldU) label, indicative of stalled or new forks, respectively. (**D**) Velocity of fork progression determined from ldU tract lengths (kb/min). (**E**) Percent of bidirectional asymmetric forks among total fork numbers. Statistical significance calculated using One-Way ANOVA followed Tukey’s Post hoc test, ****p< 0.0001, **p<0.01.

**Figure 3.**
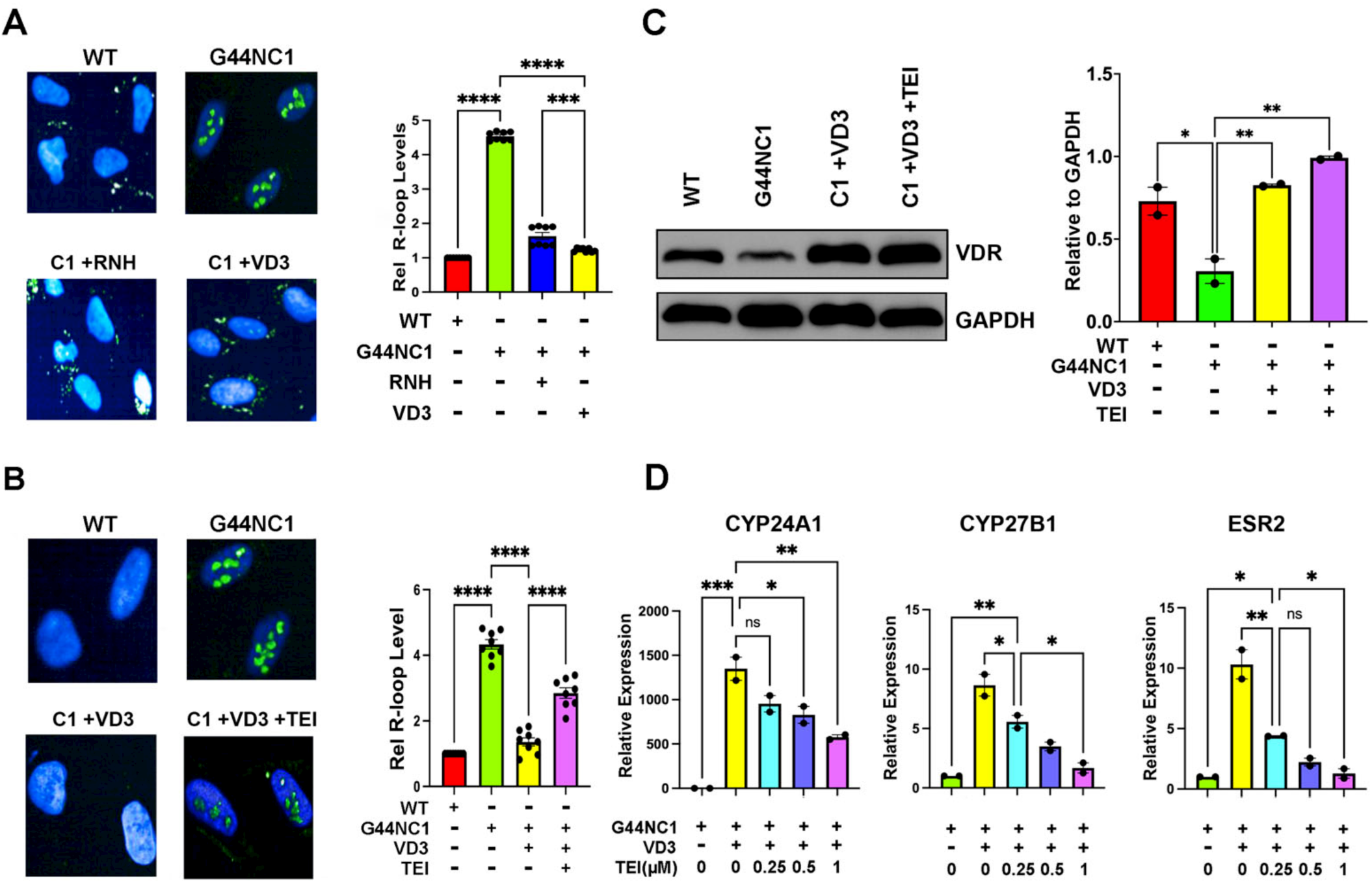
VD3 ameliorates pathogenic R-loop accrual in a VDR-dependent manner. (A) Representative images (*left*) and quantification (*right*) of R-loop levels in MED12-mutant (G44N) cells by immunocytochemical analysis using the RNA-DNA hybrid-specific antibody S9.6. [Note that one of two MED12-mutant G44N clones (Clone 1) is shown here. Analysis of Clone 2 is shown in Fig. S3]. Where indicated, cells were treated for 48h without (-) or with (+) VD3 (100nM) or transduced with RNase H1 (RNH)-expressing lentivirus prior to R-loop staining. Data are plotted as fold-change in S9.6 antibody signal intensity relative to untreated MED12 WT cells. Quantification was determined from a minimum of n=50 cells in each of 3 technical replicates from 3 independent experiments (n≥450total cells) per treatment condition. (**B**) Representative images (*left*) and quantification (*right*) of R-loop levels in MED12-mutant (G44N) cells by immunocytochemical analysis using the RNA-DNA hybrid-specific antibody S9.6. Where indicated, cells were treated for 48h without (-) or with (+) VD3 (100nM) or VD3 (100 nM) + VDR antagonist TEI-9647 (1 μM) prior to R-loop staining. Data are plotted as fold-change in S9.6 antibody signal intensity relative to untreated MED12 WT cells. Quantification was determined from a minimum of n=50 cells in each of 4 technical replicates from 2 independent experiments (n≥400 total cells) per treatment condition. (**C**) Representative immunoblot (*left)* and quantification (*right*) showing VDR expression in WT and MED12 mutant (G44N) cells. Where indicated, cells were treated for 48h without (-) or with (+) VD3 (100nM) or VD3 (100 nM) + VDR antagonist TEI-9647 (1 μM). GAPDH served as a loading control. Quantification is shown from 2 independent experiments. (**D**) Relative expression levels of VDR target genes CYP24A1, CYP27B1 and ESR were determined by RT-qPCR in the absence (-) or presence (+) of VD3 and increasing doses of TEI-9647 as indicated. Values (normalized to GAPDH) are expressed relative to that of each gene in the absence of VD3. Significance calculated using One-Way ANOVA and Tukey’s Post hoc test, ****p < 0.0001, ***p<0.001; **p<0.01 *p<0.05.

Altogether, these data indicate that VD3 ameliorates aberrant R-loop accrual and R-loop-induced replication stress in primary cells from MED12-mutant UFs and various cell-based models of MED12-mutant UFs.

VD3 has been proposed to exert its biological activities through vitamin D3 receptor (VDR)- mediated (genomic) as well as VDR-independent (nongenomic) signaling processes (48, 49). To distinguish between these possibilities, we asked if the R-loop resolving activity of VD3 was sensitive to VDR antagonism. Indeed, we found that VDR antagonist TEI-9647 (50), functionally validated herein by dose-dependent suppression of VD3-activated target genes (**Fig. 3D**), impeded the ability of VD3 to resolve pathogenic R-loops in MED12-mutant (G44N) UtSMCs (**Fig. 3B**). This finding indicates that VD3 ameliorates pathogenic R-loop accrual in a VDR-dependent manner. Notably, reduced expression of VDR in UF compared to patient-paired MM tissues has been well-documented (51–53). Similarly, we found that MED12-mutant (G44N) UtSMCs compared to their MED12 WT counterparts, expressed significantly less basal VDR protein along with comparatively higher R-loop levels, and this pattern was reversed by VD3 treatment (**Fig. 3C**). The reciprocal pattern of VDR expression and R-loop levels as a function of VD3 further supports a role for the VD3-VDR axis in R-loop suppression.

### VD3 ameliorates genomic instability driven by R-loop-induced replication stress in MED12-mutant UFs and Mediator kinase-inhibited UtSMCs

We observed a consistent 2-3-fold reduction in replication fork speeds measured in primary MED12-mutant UF cells and Mediator kinase-inhibited UtSMCs (**Figs. 1D** and **2D**). This is consistent with mild replication stress and likely explains the limited DNA double-strand break (DSB) damage observed in these cells (43), since severe replication stress instead leads to collapse of completely stalled replication forks into DSBs. Importantly, while severe replication stress limits mitotic entry of damaged DNA in checkpoint-proficient cells, mild replication stress can escape such surveillance, resulting in mitotic entry of under-replicated DNA that undergoes compensatory mitotic DNA synthesis (MiDAS) (54–57). Indeed, we observed significantly enhanced MiDAS indicative of DNA under-replication in primary cells from MED12-mutant UFs compared to those from patient-matched MM (**Fig. 4A**). Similarly, we also observed significantly enhanced levels of MiDAS in Mediator kinase-inhibited compared to control UtSMCs that was effectively reversed by ectopic overexpression of RNase H1, revealing MiDAS in this cellular context to be driven by R-loop-induced replication stress (**Fig. 4A**). Persistent under-replicated DNA can lead to chromosome nondisjunction manifested as DAPI bridges that in turn cause chromosome mis-segregation and lagging chromosome fragments, ultimately generating micronuclei (58–61). Consistent with this pathologic sequelae, DAPI bridges and micronuclei were elevated in primary MED12-mutant UF cells and Mediator kinase-inhibited UtSMCs (**Fig. 4B** and **C**) and significantly reduced by ectopic RNase H1 overexpression in the latter (**Fig. 4B** and **C**), confirming their derivation from R-loop-induced replication stress. Importantly, and consistent with its ability to inhibit R-loop-dependent replication stress, we found that VD3 suppressed elevated levels of MiDAS as well as the aberrant formation of DAPI bridges and micronuclei in primary MED12-mutant UF cells and Mediator kinase-inhibited UtSMCs (**Fig. 4A-C**). Altogether, these data indicate that VD3 suppresses chromosomal instability arising from R-loop-induced replication stress in MED12-mutant UFs.

**Figure 4.**
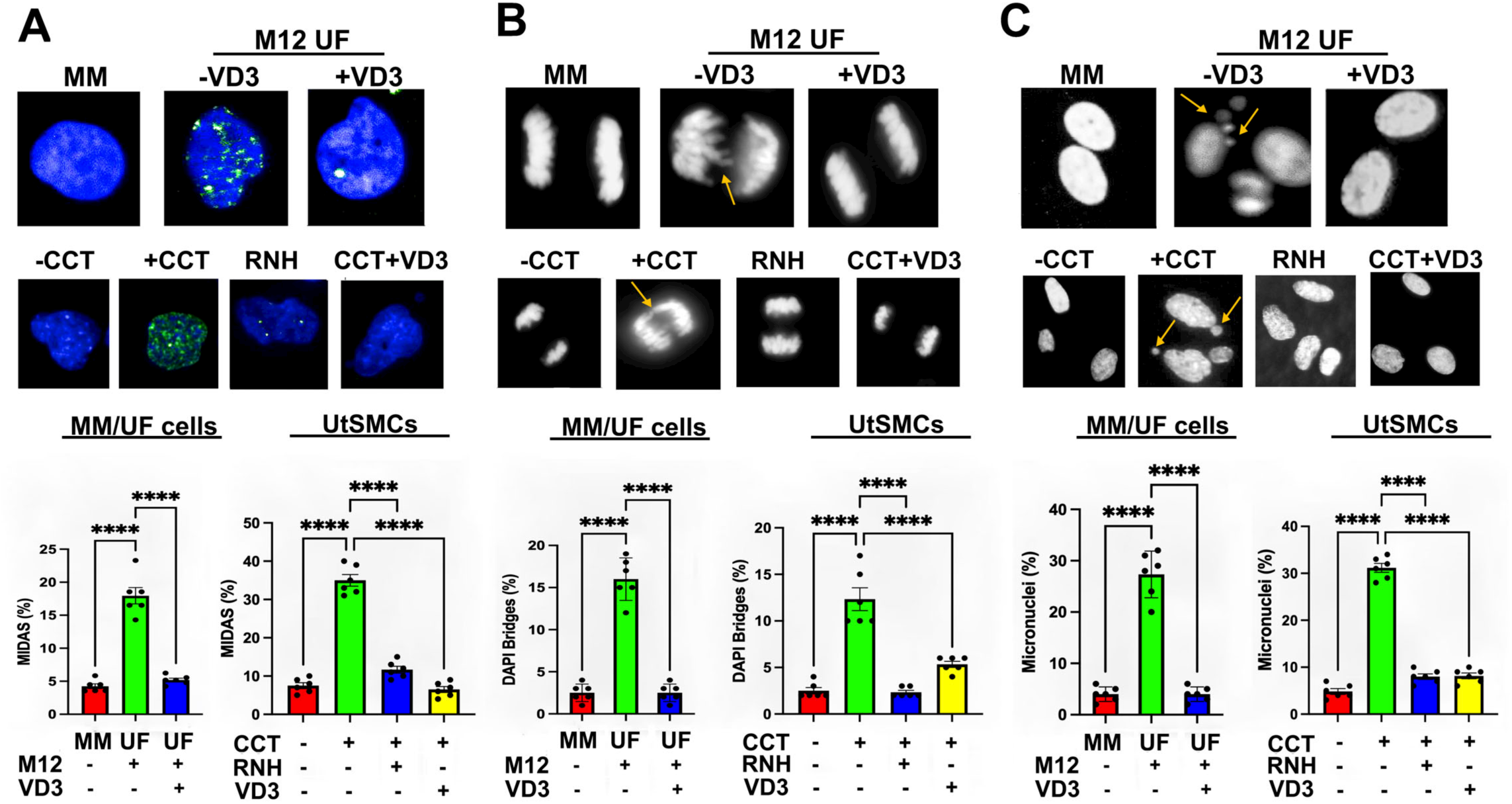
VD3 ameliorates replication-stress driven mitotic aberrations in MED12-mutant UFs and Mediator kinase-inhibited UtSMCs. Mitotic phenotypes were assessed in primary cells from patient-matched MM and MED12-mutant (M12) UFs as well as in control or RNase H1 (RNH)-overexpressing UtSMCs treated without (-) or with (+) CDK8/19 inhibitor (CCT; 100nM) for 24h. Where indicated, cells were treated with VD3 (100nM) for 48h prior to assessment of mitotic phenotypes. (**A**) Early mitotic cells were incubated with EdU for 0.5h to detect MiDAS. *Top*, representative images of MiDAS. *Bottom*, percent early mitotic cells containing EdU foci. Quantification was determined from a minimum of n=50 cells in each of 3 technical replicates from 2 independent experiments (n≥300 total cells) per treatment condition. (**B**) Anaphase cells were analyzed for DAPI bridges. *Top*, representative images of DAPI bridges (yellow arrows). *Bottom*, percent anaphase cells containing DAPI bridges. Quantification was determined from a minimum of n=50 cells in each of 3 technical replicates from 2 independent experiments (n≥300 total cells) per treatment condition. (**C**) G1 phase cells were analyzed for micronuclei formation. *Top*, representative images of micronuclei (yellow arrows). *Bottom*, percent of cells with micronuclei. Quantification was determined from a minimum of n=50 cells in each of 3 technical replicates from 2 independent experiments (n≥300 total cells) per treatment condition. Statistical significance calculated using One-Way ANOVA followed Tukey’s Post hoc test, ****p< 0.0001, ***p<0.001.

### VD3 and Doxercalciferol (DCL) suppress UF growth in a PDX mouse model of human UFs

Our observation that VD3 suppresses R-loop driven genomic instability in MED12-mutant UFs prompted us to examine its antitumor activity in a preclinical model of human UFs. Herein, we demonstrate potent efficacy of VD3 as well as Doxercalciferol (DCL), a synthetic Vitamin D2 analog, in a PDX model of human MED12-mutant UFs. The bioactive metabolite of DCL is a potent VDR agonist with similar binding affinity but a longer half-life than VD3 (62, 63). However, unlike VD3, no studies have heretofore evaluated DCL for its efficacy or ameliorative impact in women with symptomatic UFs. Accordingly, we tested DCL for its preclinical antitumor activity in our PDX model. Briefly, mice bearing MED12-mutant UF xenografts from AA patients were treated without or with VD3 or DCL for 6 weeks before sacrifice and collection of tumors for morphometric and pathological analyses. Strikingly, we observed potent dose-dependent antitumor activity of VD3, with a reduction in tumor volume approaching 70% at the highest dose administered (0.5 μg/kg/day) (**Fig. 5A** and **B**). A significant reduction in tumor volume was also observed by single-dose treatment with DCL (0.3 μg/kg/day) (**Fig. 5A and B**). Notably, at these doses, both VD3 and DCL exhibited excellent safety profiles, with no deleterious impact on liver function or calcium levels (**Fig S4A**). Histological evaluation of liver and kidney revealed no differences compared to control, except for one of four DCL-treated mice that presented calcifications (**Fig. S4B**). Further, serum AMH levels, ovarian histology, and follicular counts were unaffected compared to control (**Fig S5**). These data reveal that VD3 and DCL are well-tolerated at doses that exert significant antitumor activity.

**Figure 5.**
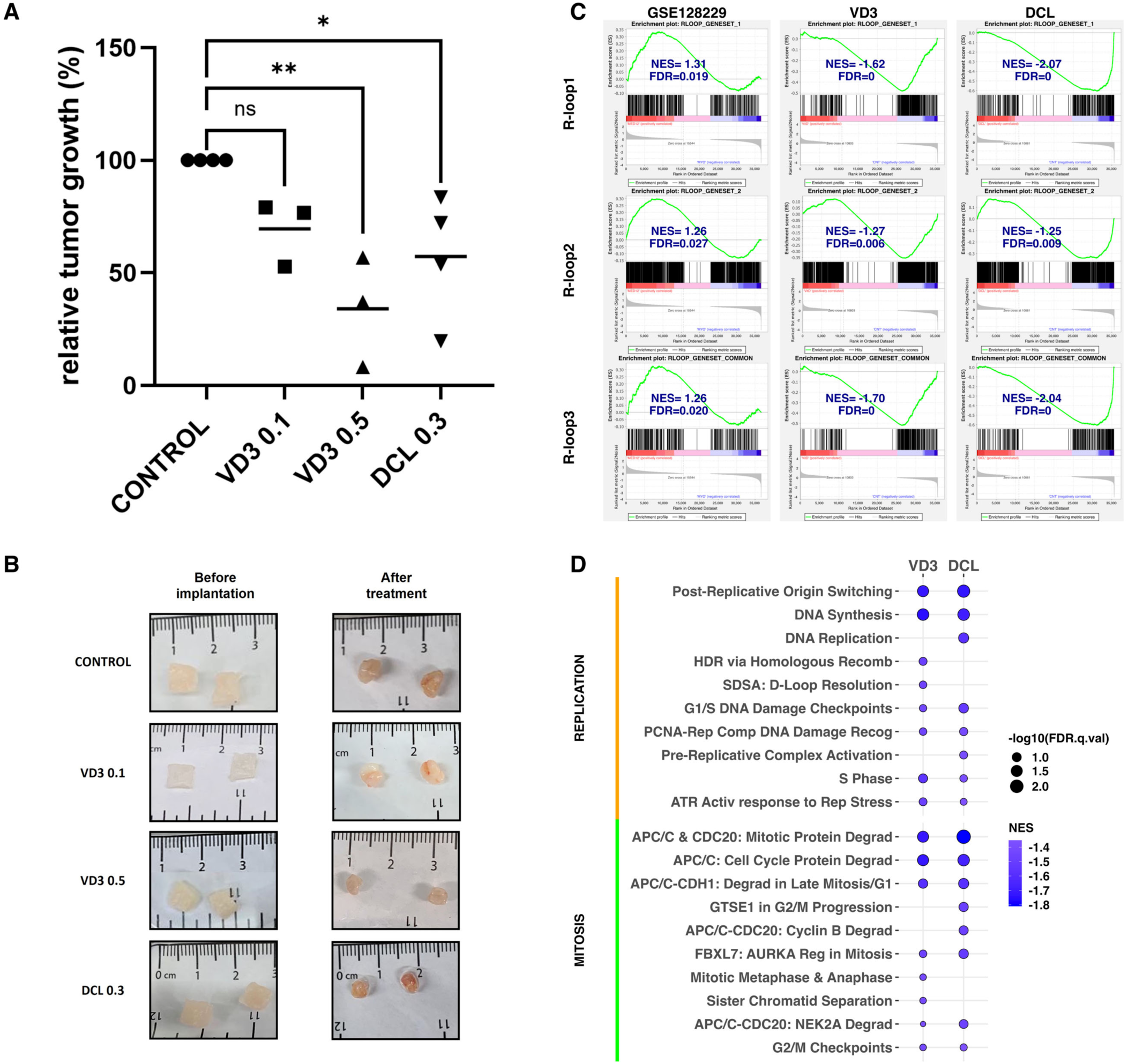
VD3 and DCL inhibit MED12-mutant UF tumor growth in a PDX mouse model. (**A**) Relative tumor growth of MED12-mutant UF xenografts treated with VD3 0.1 µg/kg/day (VD3 0.1), VD3 0.5 µg/kg/day (VD3 0.5) and DCL 0.3 µg/kg/day (DCL 0.3). Data represent mean tumor growth following 6-weeks treatment with each compound relative to its own vehicle-treated control. Significance calculated using unpaired two-tailed Student’s t-test: *p<0.05, **p<0.01. (**B**) Representative images of MED12-mutant UF xenografts before and 6 weeks after treatment with VD3 0.1, VD3 0.5, and DCL 0.3 as indicated. (**C**) GSEA plots (*left*) showing significant positive enrichment of R-loop-focused gene sets in MED12-mutant UFs compared to patient-matched MM (n=15 each) using RNA-seq data (GSE128229) reported by Moyo et al (66). GSEA plots (*middle* and *right*) showing significant negative enrichment of R-loop-focused gene sets in compound-treated (VD3 0.5 and DCL 0.3) compared to control-treated tumor xenografts. (**D**) REACTOME pathways showing significant negative enrichment in treated (VD3 0.5 and DCL 0.3) compared to control tumor xenografts. Shown are significant pathways related to DNA replication/repair and mitotic cell cycle/checkpoints.

To understand the molecular basis by which VD3 and DCL shrink MED12-mutant UFs, tumors were excised from both control and compound-treated mice at the end of the experiment and transcriptomic changes comparatively profiled by RNA-seq. We first sought to determine if VD3 (and/or DCL) reduced any tumor-associated gene signatures relevant to R-loops, as might be predicted based on our discovery that VD3 is a suppressor of pathological R-loops in MED12-mutant UFs. Because no comprehensive R-loop-specific gene set exists in the Molecular Signatures Database, we developed R-loop focused gene sets using two different sources: (1) R-loopBase (rloopbase.nju.edu) (64), a compendium of human R-loop regulators compiled by manual curation of published data prior to March 21, 2021; and (2) a recent proteomic study (absent from the R-loopBase) reliant on immunoprecipitation-mass spectrometry to identify R-loop binding proteins (65). These 2 gene sets, hereafter designated R-loop1 and R-loop2 correspond to 1185 and 364 genes, respectively. Additionally, we used the union of these 2 gene sets to identify a consensus set (hereafter R-loop3) including 280 common genes. Next, we applied gene set enrichment analysis (GSEA) to determine whether any of these *a priori* gene sets (Rloop1-3) show statistically significant concordant differences between human patient-paired MM (n=15) and MED-mutant UFs (n=15) using published RNA-seq data from Moyo et al (66). This analysis revealed significant enrichment of all three R-loop gene sets in MED12-mutant UFs compared to patient-paired MM (**Fig. 4C**; GSE128229), consistent with our findings that MED12-mutant UFs are characterized by aberrant R-loop accrual. Having confirmed these gene sets as valid surrogates for R-loop propensity in patient samples, we then applied GSEA to assess whether any of these gene sets showed statistically significant concordant differences between VD3-treated and/or DCL-treated PDX tumors compared to control-treated tumors. This analysis revealed that all 3 gene sets showed significant negative enrichment in compound-treated compared to control-treated tumors (**Fig. 4C**; VD3 and DCL), consistent with amelioration of R-loop levels by both VD3 and DCL treatment.

To further examine the impact of VD3 and DCL on biological processes that might explain their respective tumor reductive properties, we performed GSEA using the REACTOME data base. This analysis (Tables S2 and S3) revealed that VD3 and/or DCL significantly reduced canonical pathways linked to protein synthesis, cellular metabolism, and collagen deposition, consistent with the known antiproliferative and antifibrotic properties of VD3 in vitro (10–12). Notably, however, GSEA also revealed that VD3 and DCL both significantly reduced canonical pathways linked to the DNA replication stress response and DNA repair (**Fig. 4D**), as well as those involved in mitotic processes and checkpoints relevant to chromosome segregation (**Fig. 4D**), corroborating our findings herein that VD3 suppresses replication stress and chromosomal instability in primary patient-derived cells from MED12-mutant UFs. We infer that in addition to its established antiproliferative and antifibrotic properties, VD3 (and DCL) may also contribute to suppression UF tumor growth by suppression of R-loop-induced genomic instability.

## Discussion

Hypovitaminosis D is correlated with UF development, and a growing body of preclinical and clinical data support the use of VD3 as a viable therapeutic option for UF treatment. Mechanistically, VD3 has been associated with antiproliferative and antifibrotic activities in the UF setting (10–12), which likely explains, at least on part, its tumor reductive properties. Herein, we extend the role of VD3 to suppression of R-loop-induced chromosomal instability in MED12-mutant UFs, suggesting an additional basis by which VD3 may restrict tumor growth in this dominant UF subclass.

Although MED12-mutant UFs are characterized by genomic instability (27, 28, 38, 40–42), the mechanistic basis by which MED12 mutations drive this process has not heretofore been established. We found that R-loop-induced replication stress, a defining characteristic of MED12-mutant UFs (43), leads to persistent under-replicated DNA in mitosis (as evinced by ongoing MiDAS), triggering chromosomal segregation errors and formation of micronuclei, a hallmark of genomic instability. Notably, recent studies have shown that R-loops perturb replication fork progression and RNA polymerase II elongation, triggering transcription-replication conflicts (TRCs) (67, 68), that can persist through S-phase and mark sites of MiDAS repair that are susceptible to chromosomal rearrangements found in human cancers (69). Because we found that mutant MED12 promotes R-loop-induced replication stress leading to MiDAS, it is possible that chromosomal aberrations recurrently observed in MED12-mutant UFs derive from R-loop-induced TRCs that undergo attempted MiDAS repair. Furthermore, evidence supports the notion that micronuclei arising as a consequence of chromosomal segregation errors are the source of chromothripsis, a form of complex chromosomal rearrangement originally discovered in malignant tumors and, more recently, in benign tumors, including MED12-mutant UFs (38, 42). Chromothripsis refers to a process in which derivative chromosomes are generated through random re-ligation of shattered chromosome ends in a single event (70). Relevant to findings herein, chromosomal segregation defects such as those arising in MED12-mutant UFs can lead to delayed segregation of individual chromosomes and their eventual incorporation into micronuclei, which are prone to rupture and chromothriptic fragmentation by exposure to the cytosolic environment (71–73). While the precise mechanism(s) responsible have not been definitively established, proposed models nonetheless include nuclease-mediated digestion of micronuclear chromatin following rupture, condensation-dependent replication fork collapse or micro-homology-mediated annealing between stalled replication forks on micronuclear chromatin, and/or unresolved base excision repair of micronuclei-enriched R-loops (74–76). To our knowledge, this is the first demonstration that MED12-mutant UFs are prone to micronuclei formation, raising the possibility that chromothripsis in these benign tumors may derive, at least in part, from R-loop-induced replication stress and ensuing chromosomal segregation errors.

Finally, while persistent under-replicated DNA in mitosis may serve as a driver of chromosomal instability in MED12-mutant UFs, it might also represent a unique therapeutic vulnerability in this specific genetic setting. In this regard, MiDAS represents an early mitotic repair mechanism and the cell’s last chance to ensure genome integrity before cell division. Accordingly, targeted inhibition of MiDAS triggered by ongoing replications stress in MED12-mutant UFs could enhance the rate of chromosome mis-segregation and trigger tumor-specific cell death through mitotic catastrophe. Such strategies could involve targeting of factors specifically required for MiDAS in neoplastic, but not normal, cells including RAD52 and REV1 (77–80). Altogether, our findings provide new insight concerning the basis by which MED12-mutant UFs acquire chromosomal instability and suggest novel therapeutic approaches for personalized treatment of UFs stratified by driver mutation subtype.

Our studies also revealed that VD3 suppresses R-loop-induced replication stress and genomic instability in MED12-mutant UFs. Surprisingly, we discovered that this activity derives from the ability of VD3, in a manner dependent on its VD3 receptor (VDR), to suppress aberrant R-loop accrual in MED12-mutant UFs. This observation is consistent with the recent identification of VDR as a potential R-loop suppressor in an siRNA-based functional screen for drug-targetable factors that control R-loop homeostasis (81). Presently, the mechanism by which the VD3-VDR axis suppresses R-loop accumulation is unknown. It is possible that the liganded VDR could suppress pathogenic R-loop accrual indirectly through its established role as a transcriptional regulator, as for example by altering expression of known R-loop modulators. Alternatively, liganded VDR could function directly through an unanticipated role in the biochemical resolution of R-loops. Because the human VDR harbors no RNA binding motifs or nuclease/helicase domains that mark established R-loop resolving factors, it is unlikely to function independently as an R-loop processor. However, it is possible that liganded VDR could recruit R-loop resolvases in vivo, which is consistent with its established role as a hormone-dependent transcription factor known to conscript a variety of coregulator complexes (82), some of which might harbor resident R-loop processing factors.

We found that both VD3 and DCL are potent inhibitors of tumor growth in a PDX mouse model of MED12-mutant UFs. Notably, comparative RNA-seq analysis of control and compound-treated tumors revealed the latter to be negatively enriched for genesets linked to R-loops as well as biological processes linked to replication stress and mitotic progression, consistent with the ability of VD3 to suppress R-loop-induced replication stress and mitotic segregation errors in patient-derived MED12-mutant UF cells. While prior preclinical and clinical studies have both revealed VD3 to be a suppressor of UF tumor growth in vivo (11, 13–15), it’s tumor reductive properties have thus far been ascribed largely to its established antiproliferative and antifibrotic properties in vitro (10–12), with less attention given to its role in DNA repair (83, 84) and none as a regulator of R-loop homeostasis. Based on our findings herein, we suggest that VD3-imposed suppression of R-loop-induced replication stress and chromosomal instability may additionally underlie its antitumorigenic properties against MED12-mutant UFs in vivo. On the other hand, DCL has not heretofore been tested for its impact on UF tumor growth, and we now show for the first-time that this synthetic VD2 analog is indeed associated with potent antitumor activity. Clinically, DCL is indicated for the treatment of secondary hyperparathyroidism in chronic kidney disease patients, since it is activated hepatically and independent of any renal involvement (85, 86). Notably, several randomized, placebo-controlled, multicenter clinical trials have shown that DCL is effective in reducing elevated parathyroid hormone levels while restoring abnormal bone pathology without evidence of hypercalcemia or hyperphosphatemia, which is often a concern with high doses of VD3 (85, 87, 88). Accordingly, DCL or other non-hypercalcemic VD3 analogs, including paracalcitol (85, 89) or maxicalcitol (85, 86), could represent attractive alternatives to VD3 in the prevention and or treatment of UFs.

Altogether, our findings indicate that WT MED12 and VD3 both function to suppress R-loop-induced replication stress, which is unleashed by mutation of the former or reduced levels of the latter. This coordinate relationship has important implications for UF risk and treatment response, particulary for AA women who are more prone to MED12 mutation-positive tumors (30–33) and hypovitaminosis D (17–19). Thus, vitamin D-deficient women bearing MED12-mutant tumors, while at higher risk of severe disease, may also benefit most from supplemental VD3. This prediction is now amenable for direct testing through well-designed clinical trials.

## Methods

### Sex as a biological variable

Uterine fibroids are a condition observed only in females; therefore, our study exclusively examined female subjects.

### Patients and samples

This study was approved by the Institutional Review Boards of UT Health San Antonio (#20160372HU) and the University of Chicago Medical Center (#20-1414). For in vitro analyses, primary cells were isolated at UT Health San Antonio from MM and UF samples collected as fresh tissues from patients providing informed consent and undergoing hysterectomy for symptomatic UFs. Demographic and clinical pathological information for the patients is provided in Table S1. For PDX mouse model studies, UFs were collected from African American women (AA) patients providing informed consent and undergoing hysterectomy or myomectomy for symptomatic UFs at the University of Chicago Medical Center.

### Cell culture

Immortalized uterine smooth muscle cells (UtSMCs) (90) as well as hTERT and hTERT-derived MED12-mutant G44N cells (47) were cultured in DMEM/F12 (GIBCO) with 10% fetal bovine serum and 1% antibiotic-antimycotic (Invitrogen) at 5% CO2. Primary MM and UF cells were similarly cultured in DMEM/F12 (GIBCO) with 10% fetal bovine serum and 1% antibiotic-antimycotic (Invitrogen) at 5% CO2. Note that all patient-derived cells were plated immediately following isolation from intact tissues and used within 3 days for all downstream analyses to ensure retention of MED12-mutant cells that are otherwise lost within the first several passages in culture (91).

### MED12 mutation analysis

Total DNA was isolated from the myometrial and fibroid tissue using Quick Extract DNA Extraction Solution (Lucigen), according to the manufacturer’s instructions. Following total DNA extraction, the genomic region containing UF-linked MED12 mutations was PCR amplified with the following primers: MED12-F: GCCCTTTCACCTTGTTCCTT and MED12-R: TGTCCCTATAAGTCTTCCCAACC. PCR product was gel purified using QIAquick Gel Extraction Kit (Qiagen), according to manufacturer’s instructions, and Sanger Sequenced by Genewiz, (NJ, USA)

### Primary cell isolation and culture

Primary cells were isolated as described (92). Briefly, MM and UF tissues were diced manually into small pieces of <1 mm3 which were then incubated overnight (16-18h) in DMEM/F12 (GIBCO) containing 0.2% (wt/vol) collagenase (Wako), 0.05% DNase I (Invitrogen), 1% antibiotic-antimycotic mixture (Invitrogen), 10% FBS and 10mM HEPES buffer solution (Invitrogen) at 37°C on a shaker. After shaking, the material was filtered through a sterile 100-μm polyethylene mesh filter to remove undigested tissue and subsequently filtered through a 40-μm cell strainer (BD–Falcon). The filtrates were treated with ACK lysis solution for 10 min at room temperature, centrifuged and washed with 1X HBSS to remove red blood cells. Live cells were counted and seeded in 10 cm dishes (250,000cells/dish) with DMEM/F12 (GIBCO) with 10% fetal bovine serum and 1% antibiotic-antimycotic (Invitrogen) at 5% CO2. Note that patient-derived cells were plated immediately following isolation from intact tissues and used within 3 days for all downstream analyses to ensure retention of MED12-mutant cells that are otherwise lost within the first several passages in culture (91).

### RNase H1 lentivirus generation

The second-generation lentiviral system was used to generate RNase H1 WT Lentivirus. Briefly, VSV-G envelop expressing, psPAX2 packaging (a generous gift from D. Trono, Addgene plasmids #12260 and #12259), and lentiviral RNase H1-V5 transfer plasmids were transfected into HEK293T cells using X-tremeGENE HP DNA Transfection Reagent according to the manufacturer’s instructions (Millipore, Sigma). One day post-transfection, HEK293T culture medium was replaced with DMEM supplemented with 10% FBS and pen/strep antibiotics. The following day, virus was collected and concentrated at 26,000 rpm for 1 hour and 45 minutes in Optima L-100 XP Ultracentrifuge (Beckman Coulter). Thereafter, lentivirus was snap-frozen in liquid nitrogen and stored in −80 °C. To achieve transient ectopic expression of RNase H1, 70% confluent primary MED12-mutant UF cells, as well as MED12-mutant G44N cells (2 independent clonal lines named C1 and C2) were transduced with V5-tagged RNase H1 supplemented with 2μg/mL of polybrene. 48h after transduction, cells were processed for immunoblot, immunocytochemistry, and DNA Fiber analyses. To generate UtSMCs stably expressing V5-tagged RNase H1, UtSMCs (∼70% confluence) were transduced with RNase H1 lentivirus supplemented with 2μg/mL of polybrene. One day after transduction, the culture media was replaced with media containing 2μg/mL of Blasticidin (BSD) selectable marker. Stable RNase H1 expressing UtSMCs (RNase H1-UtSMCs) were selected with BSD for one week.

### Immunoblot analysis

Validation of RNase H1 overexpression in primary MED12-mutant UF cells and UtSMCs was performed by immunoblot analysis as described previously (43) using antibodies specific for the V5 epitope tag (cat # Abcam 15828). Immunoblot analysis of STAT1 in UtSMCs was performed using antibodies specific for total (SC-464; Santa Cruz Biotechnology) and phosphorylated (Ser727) (#9177; Cell Signaling Technology) STAT1. To analyse VDR expression, hTERT and MED12-mutant G44N cells were treated with VD3 (100nM) and VDR antagonist TEI-9647 (1μM, #HY-12398; MedChem Express) for 48h and immunoblot analysis was performed using antibodies specific for VDR (#12550; Cell Signaling) and GAPDH (#2118; Cell signaling) was used as the loading control.

### Immunofluorescent staining

Primary MM/UF cells, UtSMCs, hTERT, and hTERT-derived MED12-mutant G44N cells were plated in PerkinElmer cell carrier imaging 96-well plates at a density of 3000-4000 cells/well. Primary MM/UF cells were treated for 48h with either vehicle control (Ethanol) or 1α,25(OH)_2_D_3_, the bioactive form of VD3 (100nM) (Sigma-Aldrich, MI, USA), TEI-9647 (1μM), or transduced with RNase H1-expressing lentivirus prior to R-loop staining. UtSMCs were treated with vehicle control (Ethanol) or VD3 (100nM) for 48h and without or with 100nM CDK8/19 inhibitor (CCT254515; CCT) for 24h prior to R-loop analysis. For hTERT and MED12-mutant G44N cells, treatments included vehicle control or VD3 (100nM) and/or RNase H1-expressing lentiviral transduction 48h before immunofluorescent detection. Cells were fixed with 4% paraformaldehyde and processed for immunofluorescent detection as described previously (93) for R-loop detection using S9.6 antibody (ENH001, RRID:AB_2687463; Kerafast) and imaged in a single focal plane at ×40 magnification using an Operetta™ imaging system. Using Columbus™ (PerkinElmer) software, the protein intensity in the cytoplasm and nucleus were quantified. The nucleus was defined by DAPI staining (ThermoFisher Scientific) and processed for further analysis (94).

### DNA fiber analysis

For DNA fiber experiments, primary MM/UF cells were treated with control vehicle (Ethanol) or VD3 (100nM) for 48h prior to pulse labeling with 50μM IdU (Sigma-Aldrich I-7125) followed by 100μM CldU (Sigma-Aldrich C-6891) for 30min each as described previously (95). UtSMcs were treated with CDK8/19 inhibitor (CCT; 100nM) for 2h between Idu and Cldu treatments, as indicated in figure schematics. Briefly, ∼300,000 cells were embedded in agarose and DNA was prepared and then combed onto silanized coverslips using the FiberComb® Molecular Combing System (Genomic Vision). Following combing, coverslips were baked for 2 hrs at 65°C, dehydrated in ethanol (70%-90%-100%, 3 minutes each), then denatured with 0.5M NaOH + 1M NaCl for 8 minutes at room temperature. Coverslips were neutralized with PBS (3 minutes wash, 3 times), subjected to a graded ethanol series as described above and air-dried. Combed DNA was blocked with BlockAid Blocking Solution (Invitrogen B10710), followed by immunostaining with antibodies that recognize IdU (mouse anti-BrdU, BD Biosciences 347580) and CldU (rat, anti-BrdU, Abcam ab6326) for 1 hr at 37°C, washed with PBS-T, and probed with secondary antibodies (anti-mouse, Cy3, SIGMA C2181 and anti-rat, AF488, Invitrogen A11006) for 45 minutes at 37°C. followed by anti-mouse BV480 (Jackson Immuno Research 115-685-166) for 45 minutes at 37°C. Coverslips were washed in PBS, subjected to a graded ethanol series, air-dried and then mounted. Images were obtained using a 40 × oil objective on a confocal microscope Nikon Swept field. The number of stalled forks, new forks, bidirectional forks, and their lengths were measured using Image J. To calculate fork velocity, the following equation was used to convert fork length from μm to kb/min: length μm × 2.59/labeling time in min = fork velocity kb/min.

### Mitotic DNA synthesis (MiDAS) analysis

Primary MM/UF cells were treated with vehicle control (Ethanol) or VD3 (100nM) or transduced with RNase H1-expressing lentivirus 48h prior to downstream analysis for MiDAS, DAPI bridge, and micronuclei detection. UtSMCs and RNase H1-UtSMCs were treated with vehicle control (Ethanol) or VD3 (100nM) for 48h and without or with CDK8/19 inhibitor (CCT; 100nM) for 24h prior to downstream analysis for MiDAS, DAPI bridge, and micronuclei detection. For MiDAS detection in prophase/prometaphase, 60-70% confluent primary MM/UF cells, UtSMCs, and RNase H1-UtSMCs were synchronized using a double thymidine block (2mM) for 18h at 37°C. Thereafter, cells were released into prewarmed fresh media, and 9h following thymidine release, cells were treated with 20 ng/ml nocodazole for 4h. Cells were then rinsed three times (within 5 min) with pre-warmed medium (37 °C) and released into fresh cell culture media with EdU for 30min. To detect EdU, cells were first blocked with blocking buffer (3% BSA in 1x PBS containing 0.5% Triton X-100) for 30 min at RT. EdU detection was then performed using Click-IT chemistry following the manufacturer’s instructions (Click-IT EdU; Alexa fluor 594, Imaging Kits, Life Technologies). Images with EdU foci were obtained using a 40X oil objective on a confocal microscope, Nikon Swept field. Images were then processed with Image J using the same settings, and the foci were counted manually with the sample identity unknown to the counter.

### DAPI bridge detection

DAPI bridge detection was performed following a previously published protocol (96, 97). Briefly, 60-70% confluent primary MM/UF cells, UtSMCs, and RNase H1-UtSMCs were treated with demecolcine to a final concentration of 300ng/ml. After the arrest, synchronized cells were harvested by shake off and re-seeded in Poly-L-Lysine coated coverslips and were incubated for an additional 20 min to allow them to progress into anaphase for anaphase bridges analysis. Anaphase cells were first fixed with 4% paraformaldehyde for 10 min at RT followed by 3 washes with 1X PBS. Cells were then blocked with 3%BSA/PBS for 30 min at 4 °C. After blocking, cells were incubated with DAPI for 5 min followed by 3 washes with 1X PBS. Slides were mounted with Vectashield medium (Vector Laboratories) before imaging. Images DAPI bridges were obtained using a 100X oil objective on a fluorescent Nikon microscope.

### Micronuclei analysis

Briefly, 60-70% confluent primary MM/UF cells, UtSMCs, and RNase H1-UtSMCs were treated with demecolcine to a final concentration of 300ng/ml. After metaphase arrest, synchronized cells were harvested by shake off and re-seeded in Poly-L-Lysine coated coverslips and allowed to grow for an additional 2.5h to obtain G1 phase cells for micronuclei analysis. Cells were first fixed with 4% paraformaldehyde for 10 min at RT followed by 3 washes with 1X PBS. Cells were then blocked with 3%BSA/PBS for 30 min at 4 °C. After blocking, cells were incubated with DAPI for 5 min followed by 3 washes with 1X PBS. Slides were mounted with Vectashield medium (Vector Laboratories) before imaging. Images of micronuclei were obtained using a 40X oil objective on a confocal microscope, Nikon Swept field. Cell with micronuclei were counted using Image J cell counter.

### RT-quantitative PCR

Total RNA was purified using QIAzol lysis reagent and a miRNeasy Mini Kit (Qiagen, Valencia, CA). The relative abundance of each transcript was determined by RT-quantitative PCR (RT-qPCR) using SYBR Green PCR Master Mix (Bio-Rad Laboratories, Hercules, CA). The relative fold changes were calculated using the comparative cycle times method with GAPDH as an internal control on a Bio-Rad CFX384 real-time PCR detection system. Primers used for RT-qPCR were: [CYP24A Forward: 5’ CGGTGGAAACGACAGCAAA-3’; Reverse: 5’-CCATCTGAGGCGTATTATCGCT-3’], [CYP34B Forward: 5’-GGAACCCTGAACAACGTAGTC-3’; Reverse: 5’-AGTCCGAACTTGTAAAATTCCCC-3’], [ESR2 Forward: 5’TCCATCGCCAGTTATCACATCT-3’; Reverse: 5’-CTGGACCAGTAACAGGGCTG-3’], [GAPDH Forward: 5’-AACATCATCCCTGCCTCTAC-3’; Reverse: 5’-CTGCTTCACCACCTTCTTG-3’].

### Patient-derived xenograft mouse model

MED12-mutant human uterine fibroids (UF) were collected from African American women (AA) undergoing hysterectomy or myomectomy for symptomatic UFs (n=10). The collection of human tissue, retrieval and analysis of information was approved by the Institutional Review Board (IRB) of The University of Chicago Medical Center (#20-1414) and every participant provided informed consent. Female NOD-SCID mice (n=20) (Charles River Laboratories, MA, USA) were used to generate the patient-derived xenograft (PDX) mouse model. First, to maintain tumor growth, 60-day release pellets with 17ß-estradiol (0.2 mg) and progesterone (50 mg) (Innovative Research of America, FL, USA) were placed subcutaneously 1 week prior to the xenograft surgery. All the animal experiments were approved by the Institutional Animal Care and Use Committee of The University of Chicago. One week after the xenograft surgery, animals were treated for 6 weeks with the bioactive form of D3, 1α,25(OH)_2_D_3_ (Sigma-Aldrich, MI, USA) and the synthetic Vitamin D2 analog Doxercalciferol (DCL) (Hectorol; Genzyme Corp., MA, USA). According to the treatment and the dose, three different studies were conducted: (1) VD3 0.1 µg/kg/day (n=3); (2) VD3 0.5 µg/kg/day (n=3); and (3) DCL 0.3 µg/kg/day (n=4). Each study dose had its own control group, in which animals were treated with the corresponding drug vehicle. For VD3 studies, treatments were administered by micro-osmotic pumps (Alzet, CA, USA), that were subcutaneously implanted, while DCL was administered by intraperitoneal injection 3 times a week. At the end of the treatment, animals were euthanized and UF xenografts were collected. Blood, liver, kidney, and ovaries were also collected for serum determinations and histopathologic evaluation.

### Tumor growth evaluation

UF xenografts (UFx) were measured with a digital caliper before the implantation and at the end of the treatment. The volume was calculated with the use of the formula: length x width x depth x 0.52 (98). The relative tumor volume (RTV) after the treatment was calculated using the following formula: UFx final volume/UFx initial volume. The relative tumor growth (RTG) for each PDX was calculated using the formula: [(RTV of the treated UFx)/(RTV of the control UFx)] x 100.

### Safety and toxicity studies

To evaluate any significant toxicity associated with the treatments, blood samples were collected at the end of treatment and serum levels of Aspartate Aminotransferase (AST), Alanine Aminotransferase (ALT) and total Bilirubin were determined at IDEXX BioAnalytics (MA, USA). In addition, the liver and kidney were fixed, paraffin-embedded, sectioned, stained with hematoxylin and eosin (H&E) and histologically examined by pathologists who were blinded to the treatment group assignments.

### Fertility potential evaluation

To evaluate the possible effect of the treatment on mice fertility potential, serum levels of Anti-Mullerian Hormone (AMH) were determined at The Ligand Assay & Analysis Core of the Center for Research in Reproduction at the University the Virginia. Ovaries were fixed, paraffin-embedded, sectioned (5µm), and stained with H&E. The number of primordial, primary, secondary, and antral follicles were counted in three ovarian tissue sections per ovary at least 25 µm apart (each fifth section) as previously described (99).

### Library construction, RNA sequencing, and data analysis

To evaluate the effect of the VD3 0.5 and DCL 0.3 treatments on UFx gene expression, total RNA from PDX tumors was extracted using Qiazol reagent and RNeasy Micro Kit (Qiagen, CA, USA). cDNA synthesis and library generation were performed using Illumina TruSeq Stranded mRNA and sequenced in an Illumina NovaSeq 6000 (Illumina, CA, USA). The quality of reads was assessed using FastQC (http://www.bioinformatics.babraham.ac.uk/projects/fastqc, accessed on 10 Jul 2025), version 0.11.8. Subsequently, the reads were processed with Trimmomatic (100), using parameters selected based on the FastQC results. XenofilteR was employed to filter out mouse RNA stromal contamination. Briefly, reads were mapped to both the human reference genome (hg38) and the mouse reference genome (GRCm39) using STAR (101), version 2.2.1. XenofilteR (102), version 1.6, identified and removed reads that mapped better to the mouse genome, considering them as originating from the mouse host. The remaining reads were quantified with featureCounts (103), version 2.0.0, using the Gencode annotation file (104), version V41. Gene counts were pre-processed and normalized with the DESeq2 package, version 1.48.1. (105) in R. Gene set enrichment analysis (GSEA) was performed using GSEA software (106), version 4.3.3, with the REACTOME gene set downloaded from the Molecular Signatures Database. Significant pathways were determined based on 1000 permutations and an FDR threshold of < 0.25. GSEA was also performed with R-loop-focused gene sets derived from interacting genes/proteins obtained from R-loopBase (https://rloopbase.nju.edu.cn/) and Wu et al. (2021) (65). RNA-seq data from MED12-mutant UFs and matched MM were downloaded from the NCBI-GEO data repository via accession GSE128229 (66) and GSEA was performed using focused R-loop gene sets. All results were visualized using the ggplot2 R package, version 3.5.2. Raw sequencing data derived herein for PDX tumors are available through the Gene Expression Omnibus (GEO) under accession number GSE256126.

### Statistical analysis

Statistical testing was performed using Graph Pad Prism 10. One way ANOVA followed by post Hoc test was used to calculate significance. For the other experiments, including follicle counts from the PDX model, a two-tailed unpaired Student’s t-test was used. Safety data derived from the PDX model involved the use of the Kruskal-Wallis test. Significance was assumed where p-values ≤ 0.05. Asterisks represent significance in the following way: ****p ≤ 0.0001, ***p ≤ 0.001; **p ≤ 0.01; *p ≤ 0.05.

## Author Contributions

S.M., A.C., Q.Y., E.D., H.F., A.A-H., and T.G.B. conceived the study, designed experiments, and analyzed data. S.M., A.C., S.H., M.V.B., and S.K. performed experiments. S.M., A.C., E.D., H.F., A.A-H., A.K., and T.G.B analyzed the results. A.K. performed bioinformatics analysis. C.S., N.S. and R.S. obtained patient consent and performed surgeries. M.A. provided key reagents and expert insight. S.M. and T.G.B. wrote the manuscript. S.M. and A.C. were largely responsible for in vitro and in vivo studies, respectively, both of which were equally important to achieve the conclusions of the study. Based on committed effort, S.M. is listed first among the two primary co-authors. Research funding for the studies herein was acquired by A.-A-H. and T.G.B. All authors read and approved the manuscript.

## Acknowledgements

This work was supported by the *Eunice Kennedy Shriver* National Institute of Child Health and Human Development of the National Institutes of Health under award numbers R01ES028615 to A.A-H. and award numbers R01HD087417 and R01HD106285 to T.G.B. We thank the University of Chicago Genomics Facility (RRID: SCR 019196), especially Pieter W. Faber, for assistance with Illumina RNA sequencing. We acknowledge the support of the University of Chicago Animal Resources Center for assistance with animal care and regulatory compliance. Further, we thank Ms. Kyung-ah Cho (Department of Molecular Medicine, UT Health San Antonio) for expert technical assistance and tissue repository management, Dr. Srikanth Polusani (High Throughput/High Content Screening Facility, Center for Innovative Drug Discovery, UT Health San Antonio), and Dr. Exing Wang, (Director, Core Optical Imaging Facility supported by UT Health San Antonio and NIH-NCI P30 CA54174) for imaging assistance.

## Data Availability

The datasets generated during and/or analyzed during the study herein are available in the Gene Expression Omnibus (GEO) repository under accession number GSE256126.

## Ethics Approval

This study was performed in line with the principles of the Declaration of Helsinki. The Institutional Review Boards of UT Health San Antonio (#20160372HU) and the University of Chicago Medical Center (#20-1414) approved the protocols for recovery of surgical specimens.

## Consent to Participate

Informed consent was obtained from all individual participants included in the study.

