## Supplemental data for "Vitamin D3 ameliorates R-loop-induced replication stress and chromosomal instability in MED12-mutant uterine fibroids"

**Figure S1**

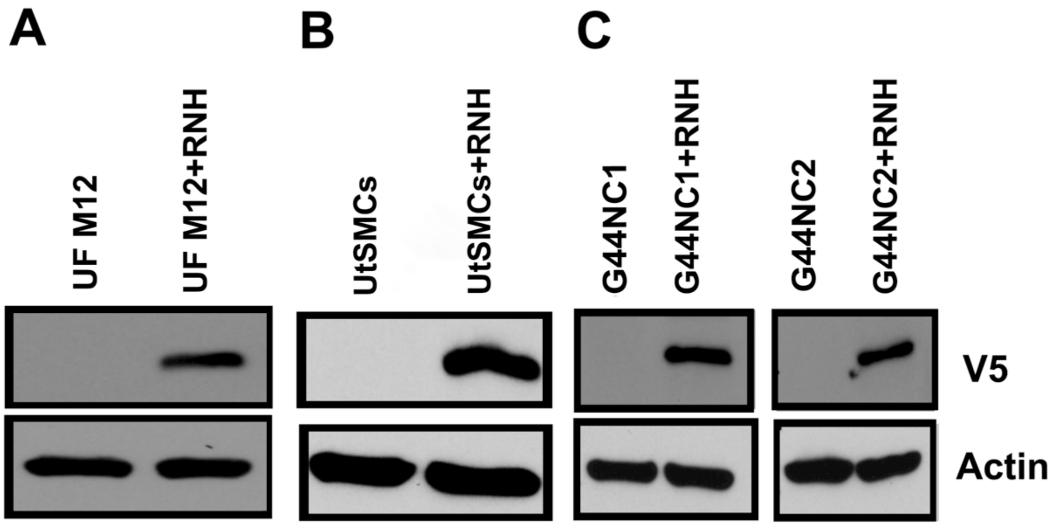


**Figure S1. Validation of ectopic RNaseH1 expression in cell culture models**. (**A)** Primary MED12-mutant **(**M12) UFs were transduced with V5-tagged RNase H1 (RNH)-expressing lentivirus in the presence 2μg/mL of polybrene. 48h post-transduction, cells were processed for immunoblot analysis using antibodies specific for the V5 epitope and β-actin, the latter of which served as an internal loading control. (**B**) Parental and V5-tagged WT RNase H1 (RNH)-expressing immortalized uterine smooth muscle cell (UtSMC) lines were processed by immunoblot analysis using antibodies specific for the V5 epitope and β-actin. (**C**) Two different CRSPR-engineered MED12-mutant G44N clones (C1 and C2) were transduced with V5-tagged RNase H1 (RNH)-expressing lentivirus in the presence 2μg/mL of polybrene. 48h post-transduction, cells were processed for immunoblot analysis using antibodies specific for the V5 epitope and β-actin.

### Figure S2

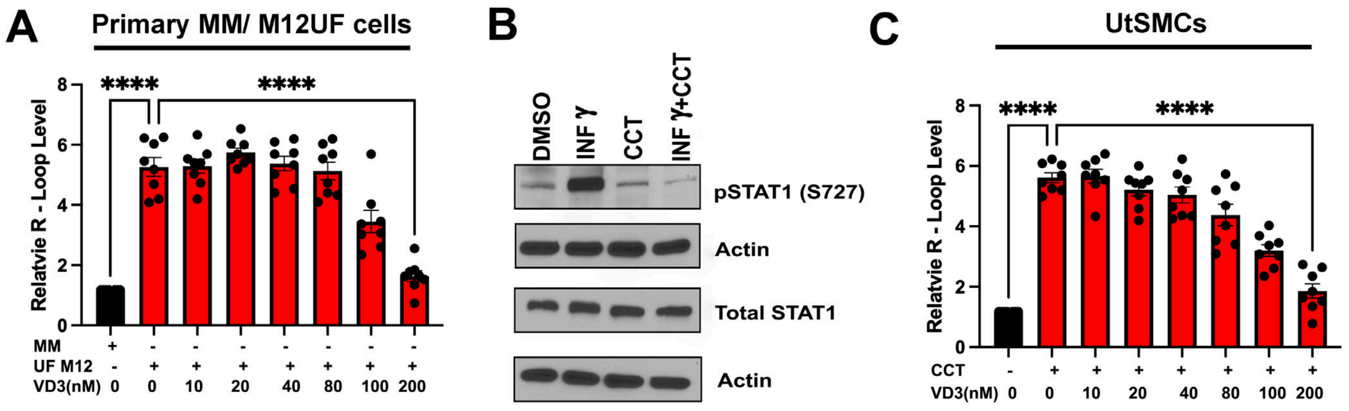


**Figure S2. VD3 dose-dependently suppresses R-loop accrual in primary MED12-mutant UFs and Mediator kinase inhibited UtSMCs.** (**A**) Quantification of R-loop levels in primary MM and MED12-mutant (M12) UFs based on immunocytochemical analysis using the RNA-DNA hybrid-specific antibody S9.6. Where indicated, cells were treated without (-) or with (+) VD3 (0-200nm). Data are plotted as fold-change in S9.6 antibody signal intensity relative to untreated MM cells. Quantification was determined from a minimum of n=50 cells in each of 4 technical replicates from 2 independent experiments ($n \geq 200$ total cells) per treatment condition. (**B**) Validation of Mediator kinase (CDK8/19) inhibition by CCT251545 in UtSMCs. As indicated, UtSMCs were pre-treated for 24h with DMSO or CCT251545 (CCT: 100nM) prior to 45min treatment without or with interferon gamma (INF$\gamma$; 10ng/ml) to stimulate Mediator kinase activity. Thereafter, treated cells were harvested and processed for immunoblot analysis using antibodies specific for phosphorylated $STAT1^{S727}$ (pSTAT1), bulk STAT1, and $\beta$-ACTIN, the latter of which served as an internal loading control. As shown by us and others, $STAT1^{S727}$ is a validated biomarker of Mediator kinase activity (1-3). (**C**) Quantification of R-loop levels in UtSMCs based on S9.6-based immunocytochemical analysis. Where indicated, UtSMCs were treated without (DMSO) or with the Mediator kinase (CDK8/19) inhibitor (CCT; 100nM) and Vit D3 (VD3; 0-200nM). Data are plotted as fold-change in S9.6 antibody signal intensity relative to untreated UtSMCs. Quantification was determined from a minimum of n=50 cells in each of 4 technical replicates from 2 independent experiments ($n \geq 200$ total cells) per treatment condition. Significance calculated using One-Way ANOVA followed Tukey's Post hoc test, ****$p < 0.0001$, *$p < 0.05$.

**Figure S3**

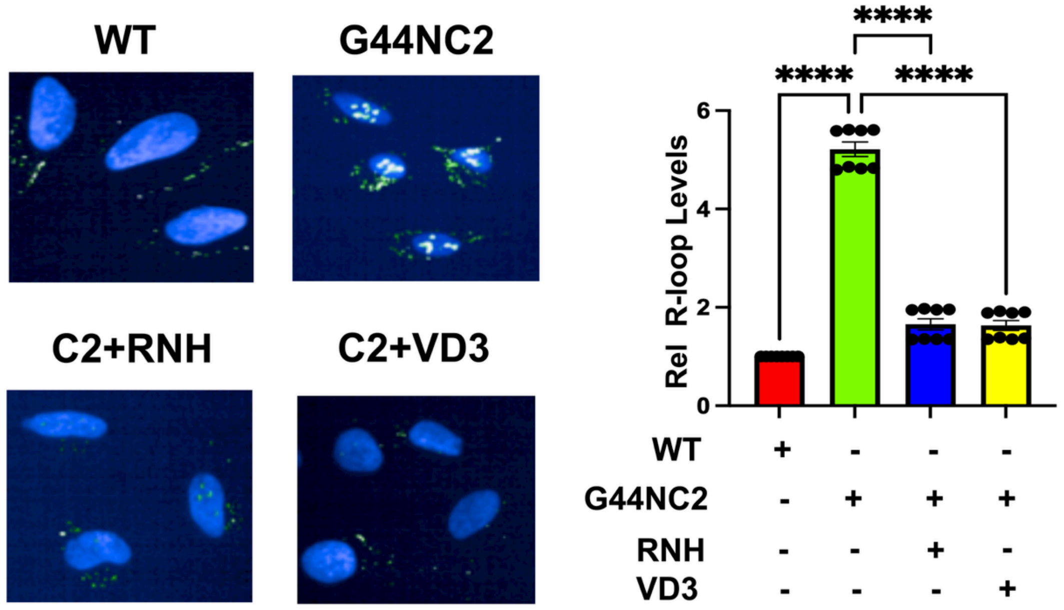


**Figure S3. VD3 suppresses R-loop accrual in MED12-mutant (G44N) hTERT cells.** Representative images (*left*) and quantification (*right*) of R-loop levels in parental MED12 WT hTERT-immortalized uterine smooth muscle (WT) cells and their derivative CRISPR-engineered MED12-mutant G44N hTERT (G44N) cells. R-loop levels were determined by immunocytochemical analysis using the RNA-DNA hybrid-specific antibody S9.6. Two different CRISPR-engineered G44N clones (C1 and C2) were analyzed. Clone 1 is shown in Figure 3. Clone 2 is shown here. Where indicated, G44NC2 cells were treated with VD3 (100nM) or transduced with RNase H1 (RNH)-expressing lentivirus. Data are plotted as fold-change in S9.6 antibody signal intensity relative to parental MED12 WT cells. Quantification was determined from a minimum of n=50 cells from each of 4 technical replicates from 2 independent experiments (n≥200 total cells) per treatment condition. Significance calculated using One-Way ANOVA followed Tukey's Post hoc test, ****p< 0.0001, ***p<0.001.

### Figure S4

**A**

| | Control n = 10 | VD3 0.1 µg/kg/day n = 3 | VD3 0.5 µg/kg/day n = 3 | DCL 0.3 µg/kg/day n = 4 | p value |
|---|---|---|---|---|---|
| **AST (U/L)** | | | | | |
| Mean ± SD | 222.4 ± 139.7 | 165.7 ± 170.0 | 236.3 ± 208.2 | 349.5 ± 215.1 | 0.4846 |
| **ALT (U/L)** | | | | | |
| Mean ± SD | 30.91 ± 10.02 | 32.33 ± 13.65 | 34.67 ± 11.59 | 30.75 ± 5.9 | 0.9384 |
| **Total Bilirubin (mg/dL)** | | | | | |
| Mean ± SD | 0.20 ± 0.04 | 0.23 ± 0.05 | 0.16 ± 0.05 | 0.22 ± 0.09 | 0.4188 |
| **Calcium (mg/dL)** | | | | | |
| Mean ± SD | 9.91 ± 0.57 | 9.53 ± 1.29 | 10.17 ± 2.17 | 11.93 ± 1.65 | 0.1865 |

**B**

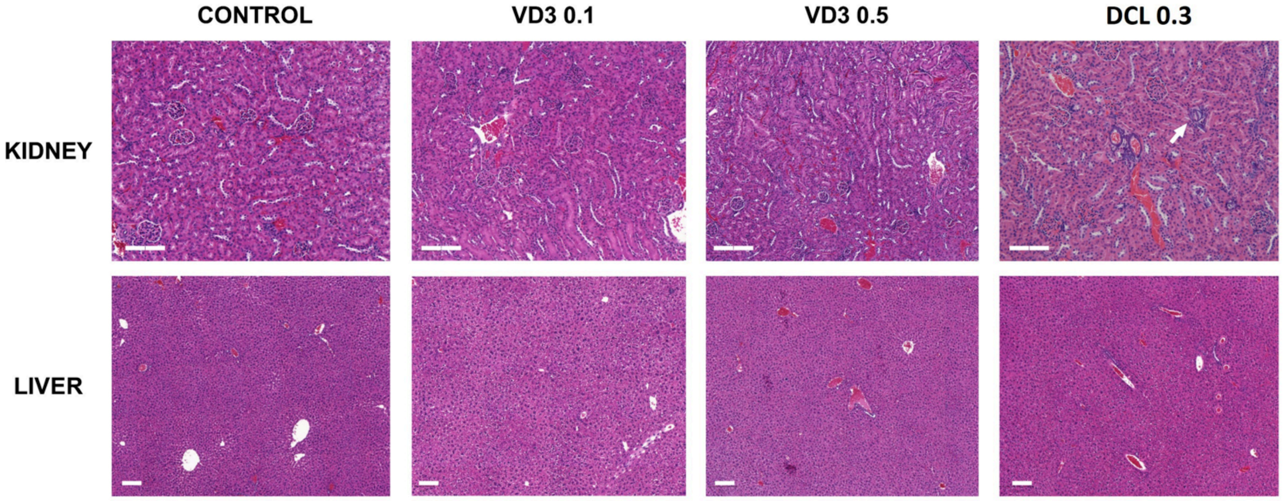


**Figure S4. Safety profiles of mice treated with Vitamin D (VD3) and Doxercalciferol (DCL). (A)** Serum levels of liver function markers Aspartate Aminotransferase (AST), Alanine Transaminase (ALT) total Bilirubin, and calcium in mice treated with vehicle (CONTROL), VD3 0.1 µg/kg/day (VD3 0.1), VD3 0.5 µg/kg/day (VD3 0.5) and DCL0.3 µg/kg/day (DCL 0.3). Values represented as mean ± standard deviation (SD). **(B)** Representative images of hematoxylin and eosin staining for histological evaluation of liver and kidney of mice from CONTROL, VD3 0.1, VD3 0.5 and DCL 0.3 groups. The white arrow points to the presence of calcifications. Scale bars: 200 µm.

**Figure S5**

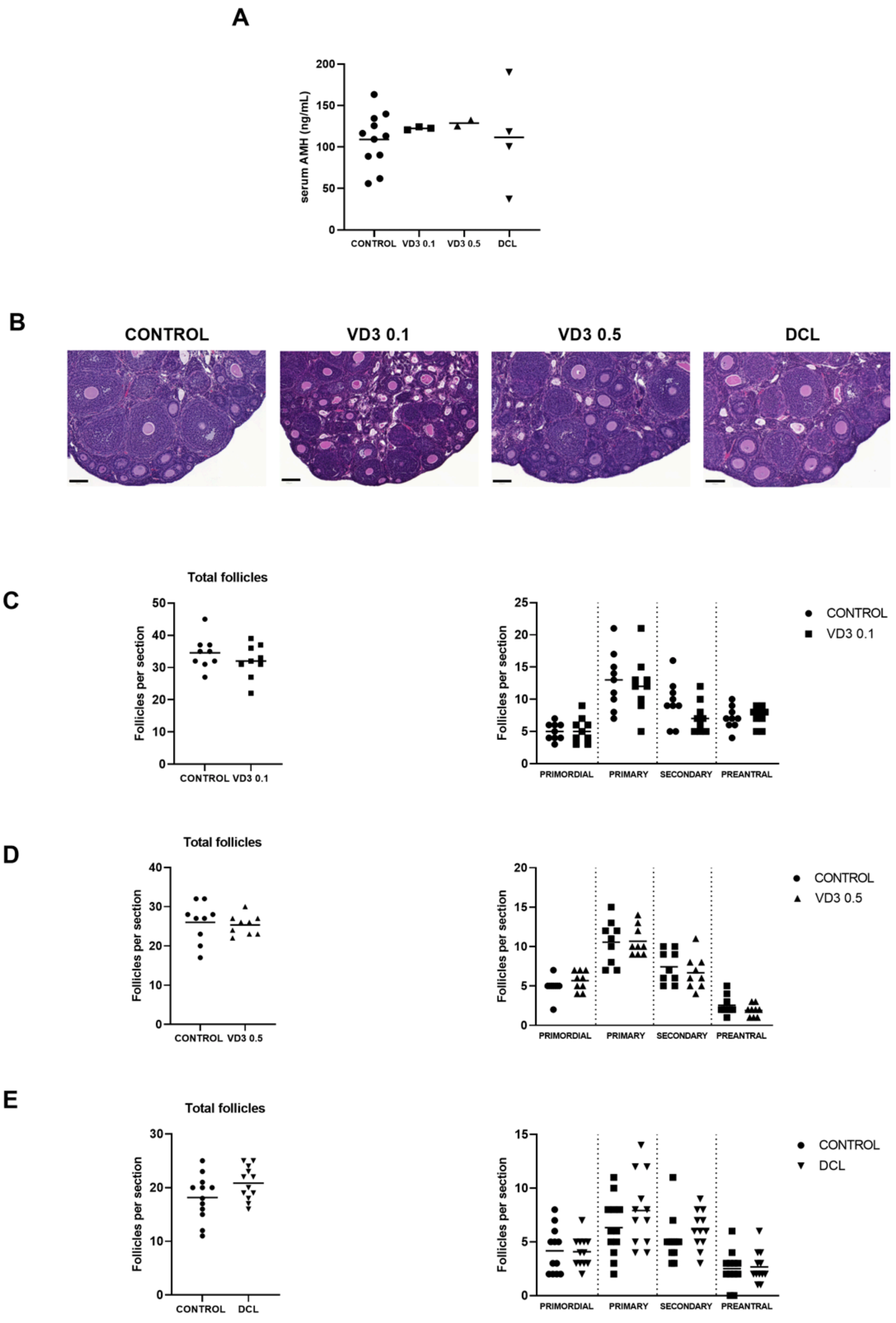


**Figure S5. Evaluation of fertility potential in mice treated with Vitamin D (VD3) and Doxercalciferol (DCL). (A)** Serum levels of Anti-Mullerian Hormone (AMH) in mice treated with vehicle (CONTROL), VD3 0.1 µg/kg/day (VD3 0.1), VD3 0.5 µg/kg/day (VD3 0.5) and DCL 0.3 µg/kg/day (DCL 0.3). Values represented as mean ± standard deviation (SD) **(B)** Representative images of hematoxylin and eosin staining for histological evaluation of mouse ovaries from CONTROL, VD3 0.1, VD3 0.5 and DCL 0.3 groups. **(C-E)** Number of total follicles and primordial, primary, secondary and preantral follicles in mice treated with VD3 0.1 **(C),** VD3 0.5 **(D)** and DCL **(E)** compared to control. Scale bars: 100 µm.

**Table S1: Information relevant to patient tissue samples used in this study**

| Sample Number | Ethnicity | Age | Solitary/Multiple fibroids | Sample ID | MM | UF M12 status |
|---|---|---|---|---|---|---|
| 1 | Non-Hispanic or Non-Latino | 29 | Multiple | 251 | Med12 WT | L36R |
| 2 | Non-Hispanic or Non-Latino | 52 | Multiple | 255 | Med12 WT | G44D |
| 3 | Non-Hispanic or Non-Latino | 49 | Multiple | 262 | Med12 WT | G44D |
| 4 | Non-Hispanic or Non-Latino | 71 | Multiple | 264 | Med12 WT | G44D |
| 5 | Unknown | 37 | Multiple | 266 | Med12 WT | G44S |
| 6 | Hispanic or Latino | 69 | Multiple | 296 | Med12 WT | G44S |
| 7 | Hispanic or Latino | 43 | Multiple | 299 | Med12 WT | G44D |
| 8 | Non-Hispanic or Non-Latino | 39 | Multiple | 314 | Med12 WT | G44V |
| 9 | Hispanic or Latino | 49 | Multiple | 338 | Med12 WT | G44S |
| 10 | Non-Hispanic or Non-Latino | 46 | Multiple | 347 | Med12 WT | G44V |
